# MUFFIN : A suite of tools for the analysis of functional sequencing data

**DOI:** 10.1101/2023.12.11.570597

**Authors:** Pierre de Langen, Benoit Ballester

## Abstract

The large diversity of functional genomic assays allows for the characterization of non-coding and coding events at the tissue level or at a single-cell resolution. However, this diversity also leads to protocol differences, widely varying sequencing depths, substantial disparities in sample sizes, and number of features. In this work, we have specifically designed a suite of tools for exploring the non-coding genome, particularly for identifying consensus peaks in peak-called assays, as well as linking non-coding genomic regions to genes and performing Gene Set Enrichment Analyses. We demonstrate that a generic but flexible count modelling approach can be utilised to compare different conditions across a broad range of genomic assay such as ENCODE H3K4Me3 ChIP-seq, scRNA-seq and TCGA ATAC-seq. Our Python package, MUFFIN, offers a suite of tools to address common issues associated with high-dimensional genomic data, such as normalisation, count transformation, dimensionality reduction, differential expression, and clustering. Additionally, our tool integrates with the popular Scanpy ecosystem and is available on Conda and at https://github.com/pdelangen/Muffin.

## Introduction

The diversity of whole-genome functionnal sequencing protocols allows us to measure a wide variety of regulatory activities across the genome. These range from transcriptomic signals, which can be detected using techniques like RNA sequencing or Cap Analysis of Gene Expression, to epigenetic signals, which can be identified with methods like ChIP-seq or ATAC-seq. The latter respectively quantify the binding of target proteins to the DNA and regions of open chromatin. The number of observations (e.g., cells or tissue samples) can also vary significantly, depending on the experimental design, from a standard two-condition triplicate protocol to thousands of observations in large-scale integrative analyses or sequencing at single-cell resolution.

These whole-genome sequencing protocols involve the mapping of sequencing reads onto a reference genome. These are then counted at specific genomic locations to effectively serve as molecular counters. The genomic locations typically originate either from reference databases (for gene-centric analyses) or are identified de-novo through peak-calling algorithms (for epigenetic or regulatory non-coding events analyses). The measured sequencing signal’s intensity can also be highly diverse, ranging from a few thousand UMIs per observation in single-cell analyses to hundreds of millions of reads in bulk sequencing. However, a common aspect across all these whole-genome sequencing protocols is that the sequencing signal per observation ultimately gets quantified through counts in a large number of features (e.g., genes or genomic peaks).

We demonstrate that a generic methodology can be employed to conduct between-observation comparisons in a wide variety of experimental designs, with low to high observation counts, and shallow to deep sequencing. This is accomplished by extending state-of-the-art approaches developed for bulk and single-cell sequencing. Notably, we build on previous statistical models and extend them to account for sequencing protocols with input or control sequencing.

Furthermore, we offer tools to facilitate work with noncoding regions of the genome. The methods presented in this paper can be easily reused and are integrated into a modular Python package, MUFFIN. Our tool is fully interoperable with the functions of the popular Scanpy tool and its ecosystem, which was initially designed for single-cell analysis, but whose tools can be adapted for broader applications.

## Methods

### Sequencing signal in arbitrary genomic features

Our framework uses raw, non-normalized read or UMI count values over genomic features. For read-based data, we provide a simple wrapper to FeatureCounts [14] to retrieve counts in BAM files located in query genomic features provided in BED, GTF, or SAF format. We support any type of genomic feature, as the user can use genes or reference maps for regulatory elements such as the ENCODE CCREs [19] or DNase hypersensitive regions catalogue [18]. For the de novo discovery of genomic elements, typically done after using a peak-calling algorithm, we provide a simple tool detailed in the next paragraph for the identification of consensus peaks across a large number of experiments. Alternatively, the user can provide their own count tables, which can be easily formatted in the standardised Python AnnData format [27] using the provided helper functions. An overview of our tools is available in Figure 1, and a detailed description of the provided tools is given in the methods below.

**Fig. 1.**
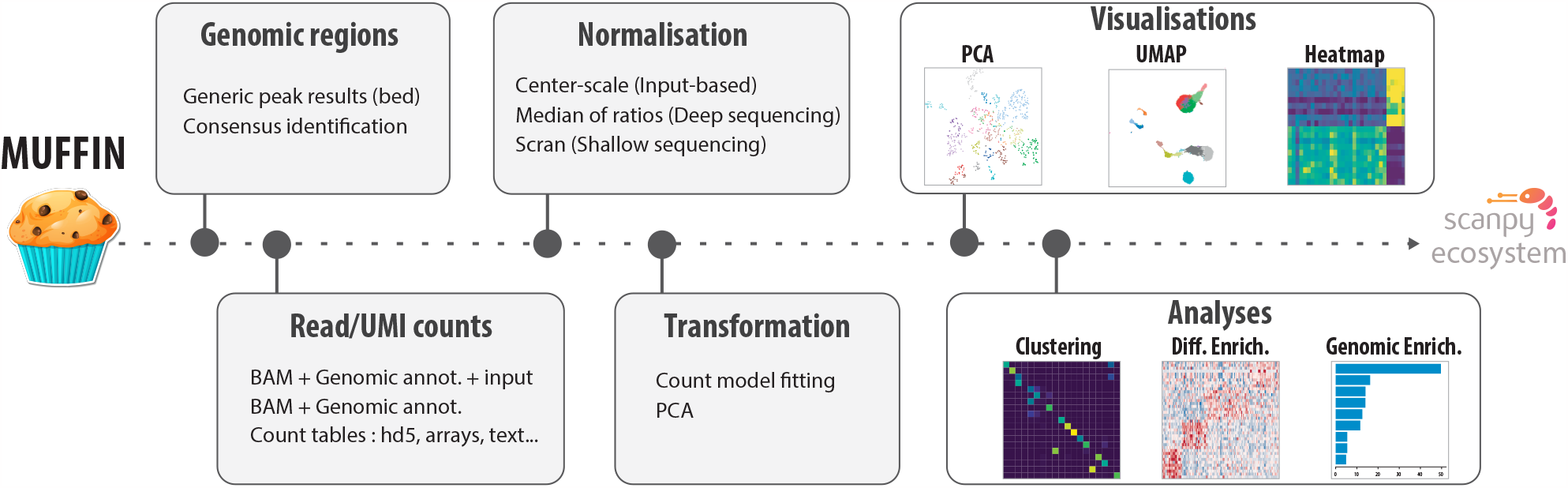
Overview of the MUFFIN suite of tools. MUFFIN covers multiple aspects of the analysis of several types of functional sequencing data, from the identification of consensus peaks, to differential expression and Gene Set Enrichment Analysis.

### High resolution consensus peak identification in large datasets

When working with sequencing data that target regulatory elements, the signal typically manifests in the form of peaks in sequencing protocols such as ChIP-seq, ATAC-seq, or CAGE. Various peak-callers have been developed to identify these signal regions on genomic tracks [23, 29]. To integrate observation-level peaks into consensus peaks, which serve as key sampling points for sequencing reads and downstream analyses, we propose a simple tool. This tool, which we used in previous work, accepts per-experiment peak-calling results (from an external tool) as input and outputs consensus peaks based on the genome-wide peak density. While identifying consensus peaks based on simple overlaps is an acceptable approach when working with a small number of experiments, it becomes problematic with a larger number of experiments. In such cases, some regions of the genome can be almost entirely covered by peaks, making the high-resolution identification of consensus peaks impossible. To merge peaks into consensus peaks, we first compute the density function of the peak summits (the single base pair genomic location with the maximum signal of the peak, we use the centre position if not available) across each chromosome. To do this, we use a kernel density estimate, employing *σ* = Median peak size / 8 as the bandwidth. We then delineate consensus peaks at each local minimum of the density function. Abnormally small artifactual consensus peaks with a size smaller than the bandwidth are discarded, as well as irreproducible peaks (consensus peaks formed by only one experiment, can be modified). Finally, we restrict the boundaries of the newly defined consensus peak to those of the farthest peaks (Figure S1). When working with stranded peak data (for example, CAGE or RAMPAGE), we simply run these steps once for each strand.

### Linking genomic regions to genes and functional annotations

Genes are much more studied and functionally annotated compared to regulatory elements of the genome. As regulatory elements are typically located near the gene they regulate, or as transcripts are frequently grouped into co-regulated clusters with similar functions, we propose tools to assist in functionally annotating regions of interest. We provide an utility to rename them according to their nearest gene, as well as a tool to perform Gene Set Enrichment Analysis based on genomic regions. We use a statistical framework that assumes the query regions are a subset of background regions (e.g., a set of differentially expressed/bound/accessible regions across two conditions is a subset of all the regions considered for differential testing). Our approach is similar to Chip-Enrich [26] and Poly-Enrich [13], which have shown that gene-wise modelling is required to reduce false discoveries. However, these two methods do not offer a model for the case where the query regions are a subset of a set of background regions, which is important as the background regions typically have a gene set bias. Following the conclusions of these two aforementioned methods, we do not use GREAT [17] as its two-tests approach saturates when querying too many regions, and as its binomial test alone is unreliable.

To assign query genomic regions to genes, we use the same heuristic as GREAT at default settings: a basal domain of 5 kb upstream and 1 kb downstream, extended in both directions up to 1 Mb or to the nearest basal domain (whichever is the closest). For each gene i, we obtain the number of genomic regions in its regulatory region, for all the background regions (*n*_*i*_) and the subset of interest (*k*_*i*_).

To compute Gene Set enrichments, we fit a Negative Binomial Generalized Linear Model per gene set s (e.g., the set of genes with the “B cell activation” GO term), which predicts the expected number of genomic regions of the subset of interest *μ*_*i,s*_ in the regulatory region of a gene i : *ln*(*μ*_*i,s*_) = *β*_0,*s*_ + *β*_1,*s*_ *× G*_*s*_ + *ln*(*E*_*i*_). Where *G*_*s*_ is equal to 1 if the studied gene belongs to the Gene Set s of interest and 0 otherwise. The term *E*_*i*_ multiplicatively corrects for the intersection bias of the background regions, with *E*_*i*_ being the expected number of hits for a particular gene: 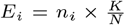, with K being the number of query regions, and N the number of background regions. We test for each gene set s whether *β*_1,*s*_ is greater than zero (i.e., whether genes in the gene set have more hits than those not in the gene set) using a Wald Test and apply the Benjamini-Hochberg FDR correction.

Additionally, to reduce redundancy between gene sets, gene sets can be clustered according to the similarities between their annotated genes. We use the leiden [25] algorithm on a kNN graph to cluster gene sets, which can be represented as a gene, gene set binary matrix of association.

### Count modelling and transformation for between observation comparisons

Following previous work [2, 15], we assume that for an observation i (e.g., sample or cell), for a variable j (e.g., genomic region or genes), the observed number of counts *K*_*i,j*_ (reads or UMIs) can be modelled with a Negative Binomial (NB) distribution with mean *μ*_*i,j*_ and overdispersion *α*_*j*_ :

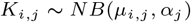

This family of distribution is discrete, heteroscedastic, skewed, and with long tails which often cause issues with approaches that assume normality. To circumvent this issue, normalizing transformations have been developed, the most popular being the log1p transform (or logCPM), which typically appears under the form :

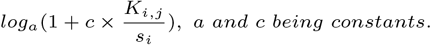

While this approach has been found to work well for large counts with few zeroes, it raises issues for small counts where it can distort the underlying distribution, especially between zeroes and small counts. Furthermore, the choice of the a and c constants is often arbitrary but does have a large effect on the balance between skewness and zero-distortion [24]. Recent work suggests using the residuals of a null (i.e constant expression) NB model that has its overdispersion parameter *α*_*j*_ being constrained to be a function of the mean [2, 7, 12]:

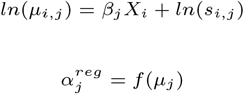

With *s*_*i,j*_ being the multiplicative normalization factor *j* for variable j and observation i; 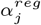 the regressed model coefficients for variable j, X the design matrix containing variables to regress out (i.e. a single intercept if there is no unwanted variables to regress). Briefly, the idea behind this approach is to model what would be the expected count distribution of a variable according to its mean expression for a constant gene/feature, then compute how much each observed count *K*_*i,j*_ is deviating from the expected fitted distribution 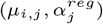 of the variable j.

In this work, we fit *f* (*μ*_*i,j*_) by randomly sampling up to 2000 variables (to speed up computations as there is no need to fit each variable to obtain a trendline), fit models using Maximum Likelihood Estimation with the Statsmodels [22] Nelder-Mead solver (without constraint on the overdispersion parameter). To obtain the regularized estimate of overdispersion we use a rolling median of the mean-sorted overdispersion values. Finally, we re-fit for each variable a new model with its regularised j using the Statsmodels IRLS solver, then compute residuals between the predicted distribution 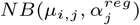 and the observed counts *K*_*i,j*_ .

The popular SCTransform approach suggests using pearson residuals, which are a linear transformation of the observed counts :

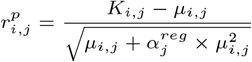

Here, we use Anscombe residuals, as implemented in Statsmodels, which are a nonlinear transformation of the observed counts, and are asymptotically following a standard normal distribution for a properly specified model.

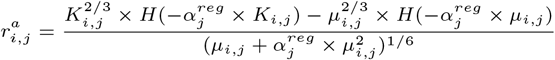

Where *H*(*x*) = *H*_2_*F*_1_(2*/*3, 1*/*3, 5*/*3, *x*), *H*_2_*F*_1_ being the Gauss Hypergeometric function.

We found Pearson residuals to be much more sensitive to outliers, and as they are still skewed and heavy-tailed due to the linear nature of the transformation, their values have to be clipped to prevent large values from driving a majority of the variance. The clipping value is a sensitive hyperparameter and has to be adjusted per dataset, with too large values being sensitive to outliers, and too small values removing biological signal. Additionally, SCTransform uses an arbitrary lower bound on the variance estimate to prevent weakly expressed genes to drive a majority of the variance. We found that Anscombe residuals do not require clipping (Figure S2), give low weights to weakly expressed features (Figure S4 A), as well as having better known theoretical statistical properties concerning normality. In our package, we implement both methods and use Anscombe residuals as the default transform.

### Between sample normalisation

We implement a few popular methods of normalisation on top of library size normalisation, such as DESeq2[15] median of ratios, Upper Quartile normalisation, and implement the scran pooling and deconvolution method [10] through a rpy2 wrapper to the scran library. The median of ratios approach is recommended for data with large counts, while the scran approach is better suited for small counts and a large number of observations. A generic method of normalisation that is independent of sample size and sequencing depth is still an area of open research. We note that our model for count distribution allows not only for a per observation multiplicative size factor but also for a per feature, per observation normalisation factor (*s*_*i,j*_). This could be exploited to use normalisation strategies that aim to correct sequence-content bias in the model. Here, we use this property to extend the model to sequencing data with input counts.

### Extending the model to sequencing data with input counts

Input sequencing is used to estimate sequence-content biases, such as PCR bias, sonication, mappability, or CNV alterations, and is used widely, for example in ChIP-seq. In these experiments, the log of the fold change (LFC), or the enrichment p-value against the input is typically used as metrics for the signal. However, the LFC does not take into account the significance of the enrichment, and on the other hand, the p-value is difficult to interpret nor displays the effect size of the enrichment; both measurements can be difficult to use for between sample comparisons. Most current methods simply ignore the amount of Input [6] and assume the sequence bias is the same across all conditions, which can be incorrect in case of differential chromatin accessibility or difference in the immunoprecipitation effiency. As the amount of input has a multiplicative effect on the observed counts, the null count model simply becomes :

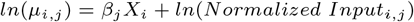

However, there is a need to normalize the input counts, and previous research has found that library size normalisation is insufficient, as the observed (ChIP) signal contains both true signal and noise reads. The signal-to-noise ratio has also been found to fluctuate strongly between experiments, mainly due to varying immunoprecipitation quality. We present a novel two-step approach, which consists of centering and then scaling the input counts. The centering step aims to find the zero of the log fold change, i.e., finds a multiplicative factor at the observation level which normalizes background regions with no signal between the ChIP and input. Here, we use the Signal Extraction Scaling [4] (SES) approach on counts sampled from random regions of the genome (10,000 per default) to estimate a centering factor *c*_*i*_. While the centering step is sufficient to perform peak calling, another scaling step is required to normalize fold changes, as samples with a more successful immunoprecipitation will have larger fold changes. We compute the per observation, median centered, signal enrichment factor *SEFM*_*i*_ as :

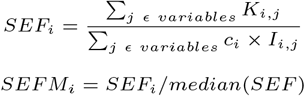

I being the input count matrix (Note that here, we do not use the counts at random genomic regions to compute *SEF*_*i*_, but the actual count matrices K and I).

Intuitively, this means that, for example, for an observation i with a two times stronger signal enrichment factor, with an observed fold change over input of 1.5 at feature j, its input should be scaled to obtain a fold change of 1.25. Formally, this translates into :

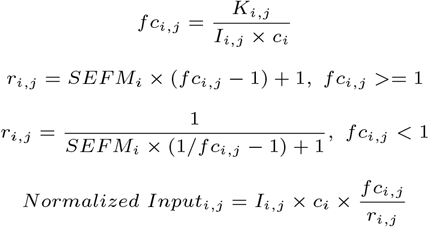

Additionally, to avoid removing features having at least a single zero as input counts (as it causes a division by 0 or taking the log of zero in the model), we replace zeroes by the mean input count for this feature (taking into account the input library size). This avoids dropping an extremely large number of variables when dealing with a large number of observations while keeping a meaningful value for the input counts.

Note that we considered ChIP-seq as an example in this paragraph, but our framework should work with experimental designs that are using a control track with an expected multiplicative effect on the resulting counts. Figure S3 highlights the effect of the two steps of our input-based normalisation.

### Differential expression

We implement DESeq2 with Log Fold Change (LFC) shrinkage [30] via an rpy2 wrapper to conduct differential expression between two conditions, as it uses a statistical model similar to ours, supporting a per feature, per observation normalisation factor. However, DESeq2 can be computationally intensive for large datasets. When working with more than 50 samples, which provides sufficient statistical power, we transition to a Welch’s t-test performed on the NB residuals, which has been shown to work well for single-cell data [2]. It should be noted, though, that the t-test offers less statistical power than DESeq2.

### Feature selection for downstream tasks

Feature selection in scRNA-seq (single-cell RNA sequencing) is a step that facilitates the elimination of a substantial portion of likely uninformative variables, i.e., those with very low expression that are mostly contaminated by technical noise, or those with ubiquitous expression, which do not provide insight into the sample/cell biology [12, 24]. Removing these can enhance the quality of downstream tasks, such as classification, clustering, or visualisation. Typically, around 2000 to 3000 genes are retained in scRNA-seq experiments, but this number is generally hand-tuned for each experiment. We use the sum of squared residuals (SSR) as a criterion for selecting variables, which serves as an indicator of the goodness of fit for the null NB model. Large residuals suggest a poor fit of the null model of constant expression, likely caused by differential expression between biological conditions that are not observed in the null model. In Figure S4 A, we show that the Anscombe residuals have a higher variance in sufficiently expressed, highly variable features.

### Optimal number of components for principal component analysis

To automatically identify the optimal number of Principal Components, we utilize Horn’s Parallel Permutation Analysis, which has proven to be one of the most effective methods to determine the number of components in factor analysis [1]. This approach involves generating row-wise permutations for each feature, computing PCA on these permuted datasets, and selecting the number of components at the threshold where the eigenvalues from the randomized dataset exceed those from the actual dataset. By default, we use the residuals as input to the PCA, and due to the computational cost of this approach, we only perform three permutations by default. This is justifiable, as the randomized eigenvalues are very stable on large matrices (Figure S4 D). To compute PCAs, we use the fast “randomized” solver from the Python sklearn library. By default, we scale PCA components to unit variance (whitening) to give an equal weight to each significant component.

### Graph Clustering

We implement the Shared Nearest Neighbour (SNN) Graph Clustering approach to identify clusters. This approach is common in single-cell RNA sequencing (scRNA-seq) analyses to identify clusters of cells without a priori on the number of clusters. To scale to a large number of points to cluster, we use an Approximate Nearest Neighbour (ANN) method to build the NN graph (python library PyNNDescent [5]). This approach avoids the quadratic time complexity of building exact nearest neighbours, can use any metric and runs in an almost linear time complexity. By default, we use the PCA representation with pearson correlation as the metric to build the NN graph. In the SNN graph, vertices are weighted by the number of shared nearest neighbours between the two nodes. To identify communities in the SNN graph, we used the Leiden graph clustering algorithm [25] implemented in the Python Leidenalg library.

### Data visualisation

For visualising the similarity among observations in datasets with a large number of features and observations, dimensionality reduction methods such as UMAP (Uniform Manifold Approximation and Projection) or t-SNE[16] (t-Distributed Stochastic Neighbor Embedding) have become standard, particularly for the analysis of single-cell data. Here, we provide a wrapper for UMAP, which is, by default, performed on the PCA representation. By default, Pearson correlation is used as the metric when there are more than 10 input dimensions; otherwise, the Euclidean distance is used.

### Analysis of benchmark datasets

We manually retrieved matching peak calling BED files, ChIP BAM files, and Input BAM files from ENCODE [19] (see Supplementary Tables). We selected only those H3K4Me3 ChIP-seq experiments that were mapped onto hg38, and removed samples with any audit errors. In cases where multiple replicates of the input file were present, we retained only the most deeply sequenced one and used it across all biological replicates. We used a coarse annotation (the same as in the ENCODE data portal) instead of the detailed cell types as there is a large number of precise cell types with no replicates. The list of genes associated with Gene Ontology terms was retrieved from the g:ProfileR website[21].

We obtained the filtered 10k PBMC scRNA-seq count tables from 10X Genomics (https://www.10xgenomics.com/resources/datasets/10k-human-pbmcs-3-ht-v3-1-chromium-x-3-1-high). We also analyzed the dataset using Seurat with SCTransform V2 as a reference, with the default settings from its tutorial vignette. We compared the similarity between our method and SCTransform using the jaccard index of similarities between clusters, and using Adjusted Mutual Information (AMI) to compare similarities across all clusters.

TCGA ATAC-seq[3] count tables were obtained from the NIH (https://gdc.cancer.gov/about-data/publications/ATACseq-AWG). The list of candidate genes associated with specific cancer hallmarks was obtained from the CHG database[28].

All analyses were conducted using the default settings, with the exception of the normalisation method: for the single-cell dataset, we employed scran normalisation; for the TCGA ATAC-seq dataset, we applied DESeq’s median of ratios; and for the ChIP-seq dataset, we used our custom normalisation method. Furthermore, we utilised logistic regression via Scanpy to identify markers across multiple classes in both the single-cell and the ATAC datasets.

## Results

### MUFFIN Suite: applications and validation in diverse biological contexts

Here we introduce the comprehensive MUFFIN suite, encompassing a diverse array of tools tailored for analyses across a spectrum of experimental designs, accommodating varying observation counts and sequencing depths. The origin of this suite can be traced back to the creation and analysis of an RNA Polymerase II Atlas, involving the development and use of various tools in the context of 900 ChIP-seq experiments and approximately 28,000 RNA-seq samples [11]. The suite’s functionalities include the analysis of sequencing signals, in the form of raw non-normalized read or UMI counted over a range of genomic features such as genes, or reference maps for regulatory elements like ENCODE CCREs or DNase hypersensitive regions. Additionally, the suite offers high-resolution consensus peak identification for large datasets, integrating observation-level peaks into consensus peaks—a crucial feature when dealing with a larger number of experiments. Tools are also provided for linking genomic regions to genes and functional annotations, including utilities for renaming regions based on their nearest gene and performing Gene Set Enrichment Analysis. Furthermore, the suite encompasses a variety of tools for functional sequencing analyses, covering count modeling for between-observation comparisons, differential expression analysis, feature selection for downstream tasks, determination of the optimal number of components for principal component analysis, graph clustering, and data visualization (1). A detailed description of the tools is available in the Methods. To validate the suite’s efficacy, it was tested in three largely different datasets : i) the ENCODE Epigenetic Atlas of Immune Cells using H3K4Me3 ChIP-seq, ii) 10,000 PBMCs sequenced with 10X scRNA-seq, and iii) 400 cancer samples sequenced via ATAC-seq originating from TCGA. These applications showcase the versatility and robustness of the MUFFIN suite across diverse biological and technical contexts.

### ENCODE epigenetic atlas of Immune cells through H3K4Me3 ChIP-seq

In order to evaluate the general applicability of our proposed methodology, we utilised the H3K4Me3 ChIP-seq dataset from the comprehensive ENCODE epigenetic atlas of immune cells. H3K4me3, an epigenetic modification known for marking active promoters, is a crucial mediator of transcriptional activity and plays a significant role in cellular differentiation and function. Here, multiple difficulties are linked to this dataset : there are no reference regions to sample sequencing signals from, the sample sizes are rather small, and the sequencing protocol uses a control track.

Our first task was to identify which regions to consider for quantifying H3K4Me3 occupancy. For this purpose, we employed our density-based approach to identify consensus peaks from per-experiment peak calling results. Our approach identifies the consensus peaks at high resolution in dense regions, and correlates with the Transcription Factor binding density from the external ReMap database[8] (Figure 2A). Subsequently, we counted reads in immuno-precipitated and input sample pairs at consensus peak locations. Ultimately, the counts are normalised using our centering and scaling approach (Methods), transformed using Anscombe residuals, and can be processed through a visualisation or dimensionality reduction routine. As shown in Figure 2 B, a heatmap view shows a clear distinction between immune cell types and reveals distinct H3K4Me3 occupancy across different cell types.

**Fig. 2.**
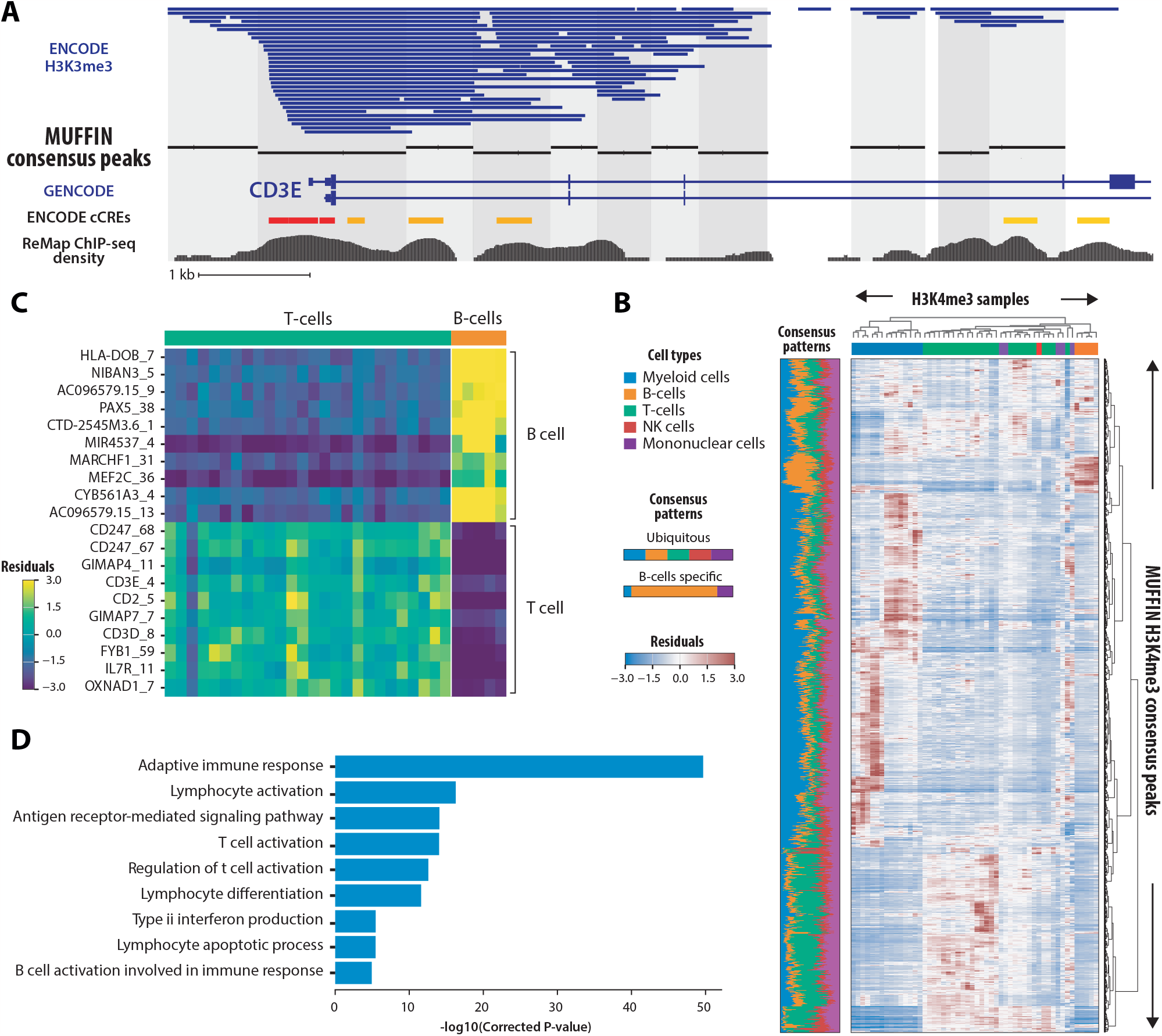
Analysis of the ENCODE epigenetic atlas of Immune cells through H3K4Me3 ChIP-seq. **A**. Result of the consensus peak identification in a 10kb region around the CD3E gene. Top to bottom rows indicates: ReMap Transcription Factor binding density, Consensus peaks, Peaks from all experiments aggregated, GENCODE V43 annotation. The alternating colors highlight the coverage of each consensus peak. **B**. Heatmap of the Anscombe residuals. Rows correspond to samples and columns to consensus peaks. Bottom panel indicates the weighted proportion of the total Sum of Squared Residuals carried by each class. **C**. Heatmap of the Anscombe residuals, for the top 10 most differentially marked consensus peaks (each renamed according to its nearest gene) in either B-Cells or T-Cells. **D**. Clustered GO terms enrichments of genes near differentially marked consensus peaks between B-Cells and T-Cells. Terms are clustered by gene similarity and only the term with the strongest enrichment is displayed.

Next, we sought to identify regions differentially marked by H3K4Me3 between B-cells (n=5) and T-cells (n=26). We found that the regions most differentially bound in B-cells are indeed located near B-cell marker genes such as HLA-DOB, which is expressed in Antigen Presenting Cells, or PAX5, a transcription factor involved in B-Cell differentiation. Conversely, for T-cells, we also retrieved T-cell markers like CD247 (T-cell surface glycoprotein CD3 zeta chain) or CD3E (CD3 Epsilon Subunit Of T-Cell Receptor Complex). Using our tool of Genomic Regions Enrichment Analysis, we find that these differentially occupied regions are located near genes annotated with immune and lymphocyte-related Gene Ontology terms (Figure 2 C-D). As a whole, this confirms that our methods are able to analyze ChIP-seq data and are likely to work well with other similar datasets that are using control sequencing.

### Cell-type clustering of 10,000 PBMCs sequenced by 10X scRNA-seq

After investigating the differential occupancy of H3K4Me3 across various immune cell types, we sought to evaluate the robustness and versatility of our methodology using another widely adopted dataset. Hence, we turned our focus to a single-cell RNA-sequencing (scRNA-seq) dataset, which offers a distinct set of challenges due to the large number of observations, inherent sparsity, shallow sequencing depth and high dimensionality of the data. The 10X Genomics scRNA-seq dataset of Peripheral Blood Mononuclear Cells (PBMCs), a mixed population of various immune cells, is often used as a benchmark case study. The PBMC dataset encompasses numerous immune cell types, allowing us to assess the efficacy of our approach in identifying and distinguishing between different cellular phenotypes.

Here, count quantification per gene, per cell has already been performed. We used UMAP to visualise the transcriptomic similarities between individual cells and applied graph clustering to pinpoint cell types (Figure 3 A). We clearly identify canonical cell types, such as Monocytes, B-Cells, T-Cells or platelets. Their markers, identified via the coefficients of a logistic regression, are also coherent (Figure 3 B), with canonical markers such as CD8B for CD8+ T-Cells, or GP1BB for platelets. Furthermore, we are able to identify cell types at a fine resolution, highlighting clusters of cells with similar transcriptomes, but with a few very specific markers, especially in the B-Cell clusters and the monocyte clusters. Overall, our methodology was able to provide clusters similar to those obtained with the reference SCTransform method (Figure 3 C). Together, these results show that our approaches are also well-suited for the analysis of datasets with low sequencing depth that are typically found in single-cell sequencing.

**Fig. 3.**
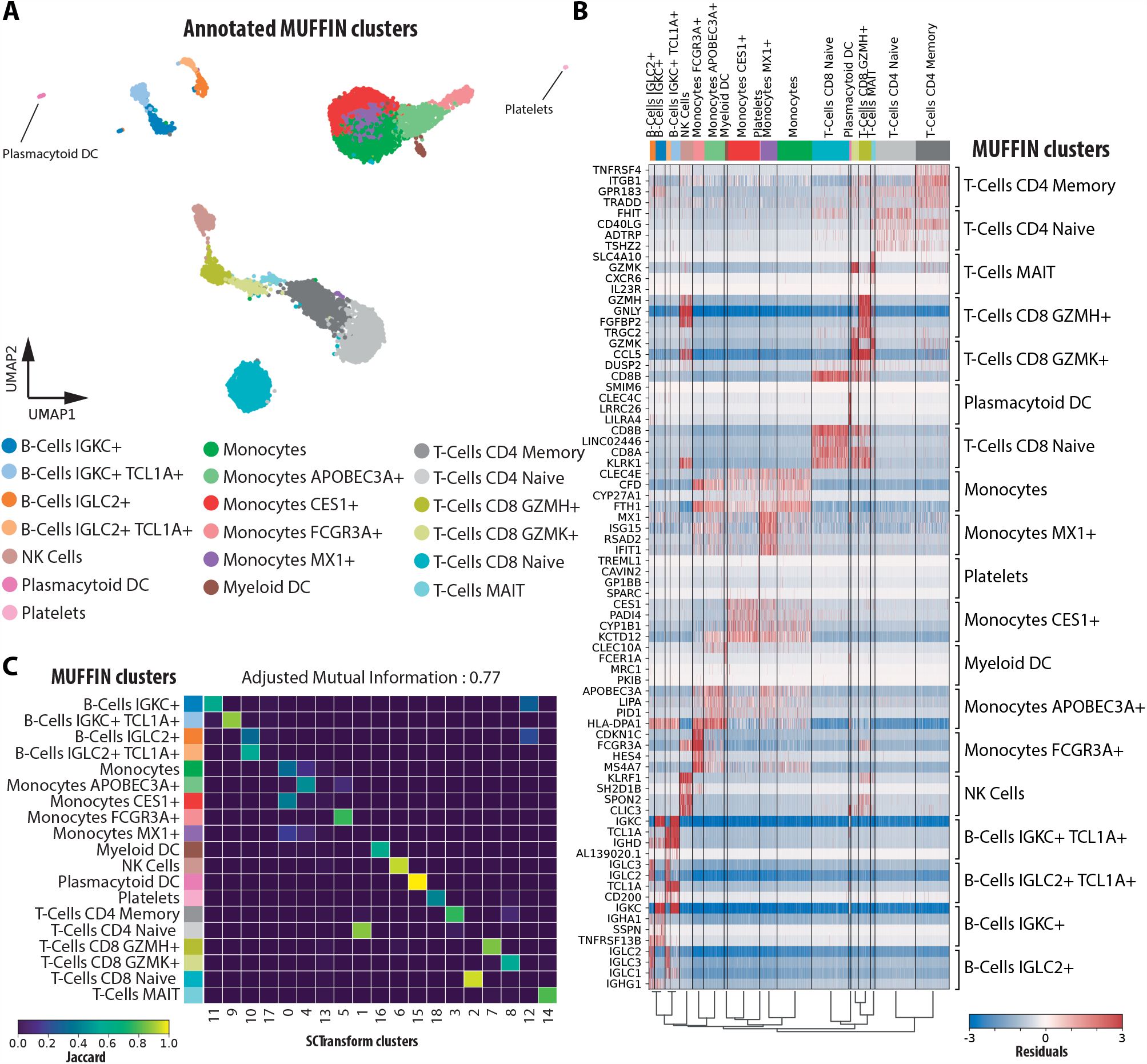
Analysis of a 10k PBMCs single-cell RNA-seq dataset. **A**. UMAP visualisation of the annotated clustered cells. **B**. Heatmap of the Anscombe residuals, per cell, for the 4 best markers of each cluster, identified via the coefficients of a multivariate, multi-class logistic regression. **C**. Heatmap of the Jaccard indexes between clusters identified via SCTransform and MUFFIN.

### The Cancer Genome Atlas open chromatin landscape of cancers via ATAC-seq

Finally, we studied the open chromatin landscape in cancers using Assay for Transposase-Accessible Chromatin using sequencing (ATAC-seq) in tissues. ATAC-seq is a technique used to assess the chromatin accessibility, thereby providing insights into the regulatory regions active in a particular cell type, tissue or condition. This sequencing protocol typically generates deeply sequenced datasets with an extremely large number of features.

The read count table per ATAC peak, per sample was already provided, but a consensus peak identification as performed with the ChIP-seq dataset could have been performed. In Figure 4 A, using UMAP, we can clearly distinguish the different cancer types, and even distinguish between different breast cancer molecular subtypes. When focusing solely on breast cancer samples, we can identify markers for each subtype (Figure 4 B). While Luminal A and B subtypes do not exhibit strongly characteristic signatures, the Basal and Her2 subtypes display characteristic regions of open chromatin. Notably, in the Her2 subtype, an open chromatin region located near the main marker of this cancer, ERBB2 (or Her2) can be identified. Using our tool of Genomic Regions Enrichment Analysis, we observe that characteristic open chromatin regions for each breast cancer subtype locate near cancer hallmark genes (Figure 4 C). Notably, the Her2+ breast cancer subtype is known to have elevated levels of Vascular Endothelial Growth Factor (VEGF) gene expression, inducing angiogenesis [9, 20].

**Fig. 4.**
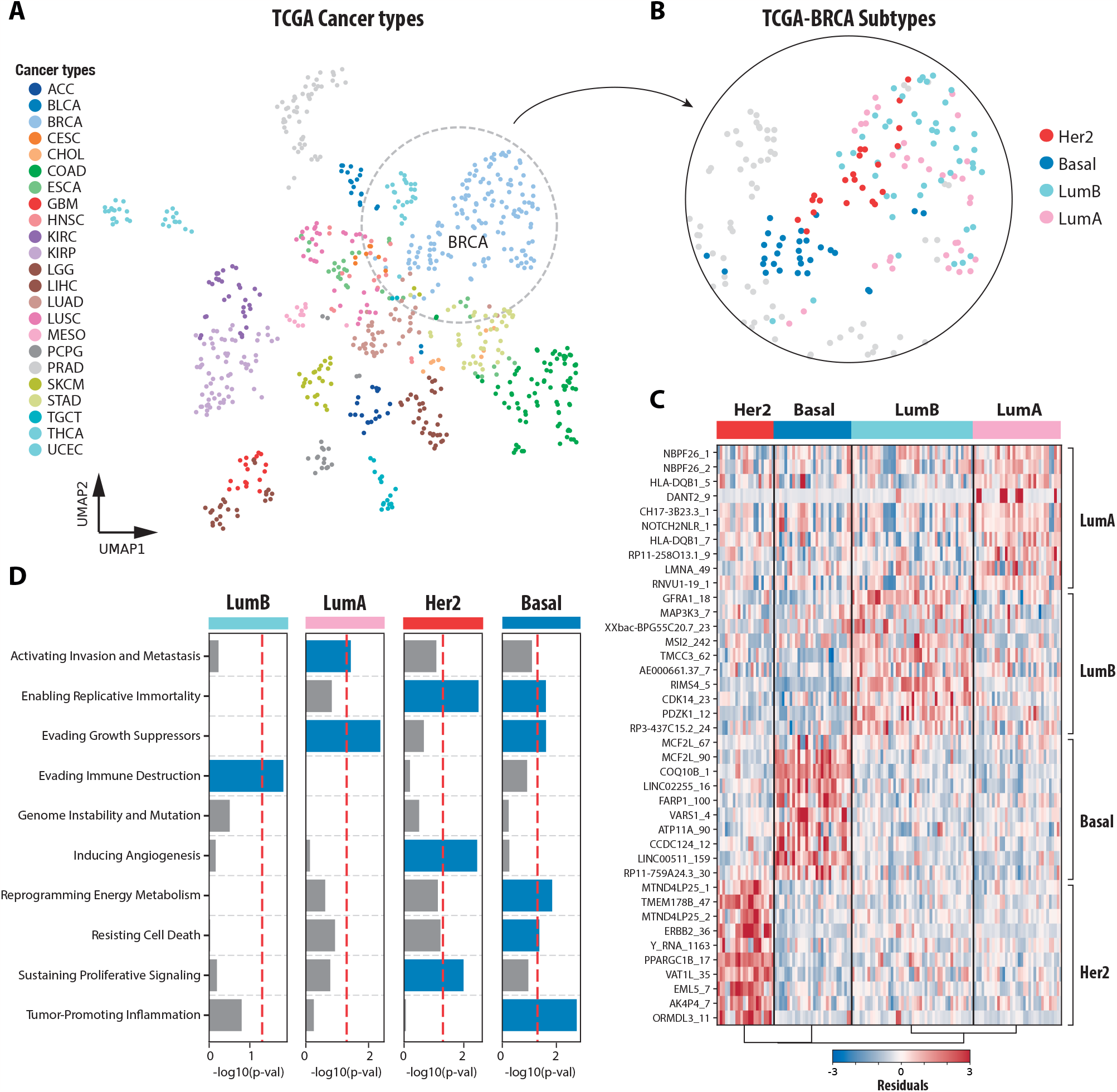
Analysis of cancer samples sequenced with ATAC-seq. **A**. UMAP visualisation of each tissue sample, annotated by cancer type, as well as subtype for the breast cancer samples. **B**. Heatmap of the Anscombe residuals, per sample, for the 10 best markers of each breast cancer subtype, identified via the coefficients of a multivariate, multi-class logistic regression. **C**. Enrichment (-log10(p-value)) in cancer hallmarks for genes nearby the 5% most discriminative ATAC peaks for each breast cancer subtype, identified via the coefficients of a multivariate, multi-class logistic regression.

## Discussion

In conclusion, our proposed framework, MUFFIN, represents a valuable and generic toolset for the processing and analysis of count data derived from a variety of high-throughput sequencing experiments. We have demonstrated its performance across diverse datasets, from epigenetic to transcriptomic, and from bulk to single-cell resolution. The underlying model is highly flexible and can adapt to a wide range of experimental designs, and relies on already well-tested, state-of-art statistical approaches.

One of MUFFIN’s key strengths lies in its ability to address multiple facets of count-based genomic assays in a generic way with minimal hand-tuning : normalization, count transformation, feature selection, dimensionality reduction, visualization, differential expression, and clustering. Additionally, it offers specialized tools for analyzing data that do not rely on gene annotations. This includes a tool for identifying consensus peaks at a high resolution, and another one for performing functional enrichment of genes located near genomic regions of interest. Furthermore, MUFFIN seamlessly integrates in the existing Python Scanpy ecosystem and its diverse range of tools. The modular architecture of MUFFIN could also allow future packages to be built on top of it.

However, it’s worth noting a potential limitation of MUFFIN, which is its compatibility with extremely large, sparse datasets. As its count transformation does not support sparse data formats, this may restrict its use in situations where memory capacity is a significant concern, such as in the analysis of extremely large integrative single-cell datasets consisting of millions of cells.

In summary, MUFFIN offers generic tools to analyze high-throughput sequencing count data and complements the existing tools available in Scanpy and the Python ecosystem. We anticipate that it will be of strong interest for bioinformaticians working with functional genomic data, at the tissue or single-cell level.

## Author contributions statement

P.D.L. took the lead in developing the methods, coding, and analyzing the datasets. Both P.D.L. and B.B. contributed to the writing and reviewing of the manuscript. All authors edited and approved the article.

## Acknowledgements

We would like to thank Dr. Lionel Spinelli for engaging in discussions that have contributed positively to our work.

This work was supported with ; PhD Fellowship to P.D.L. from the French Ministry of Higher Education and Research (MESR); Institut National de la Santé et de la Recherche Médicale (INSERM); The Core Cluster of the Institut Français de Bioinformatique (IFB) (ANR-11-INBS-0013) for granting access to its high performance computing resources. The results shown here are based upon data generated by the TCGA Research Network, the ENCODE Consortium and the ENCODE production laboratories.

---

**Fig. S1.**
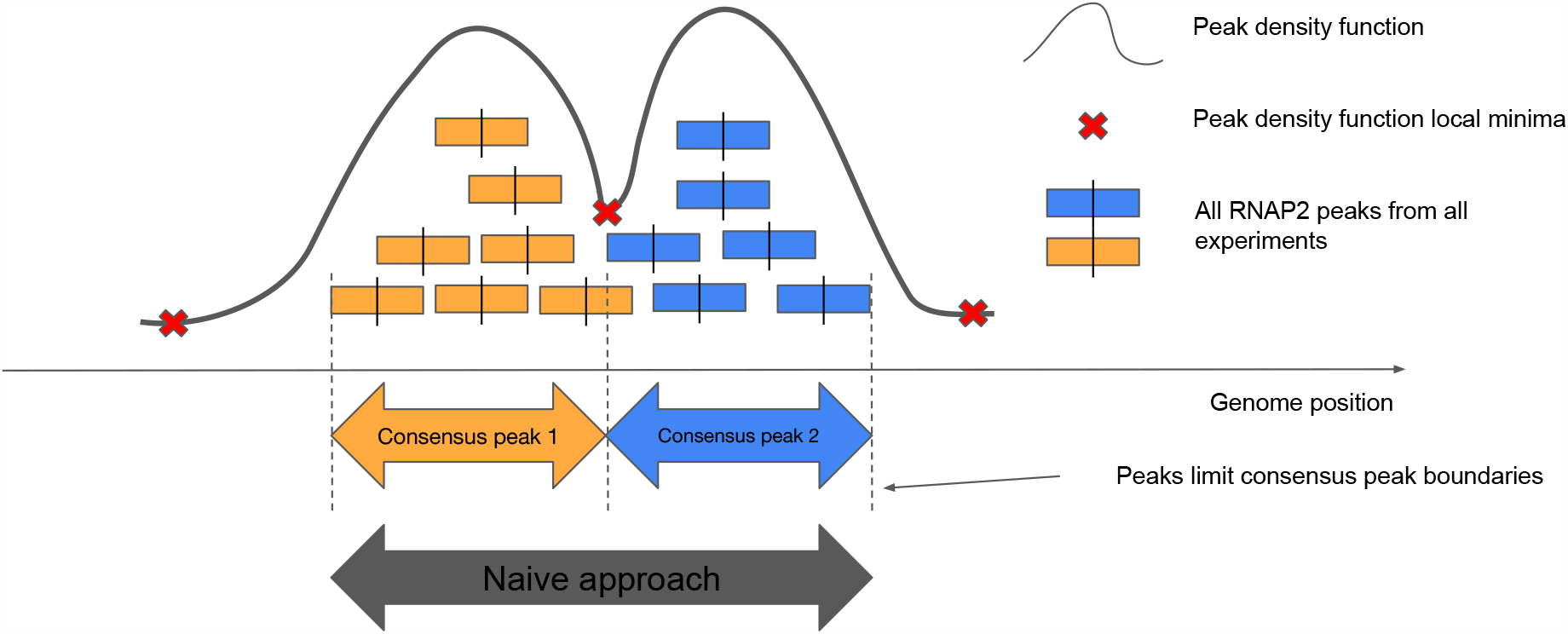
Identification of consensus peaks. Identification of consensus peaks via a peak density-based approach. Naive approach refers to a simple merge on overlap.

**Fig. S2.**
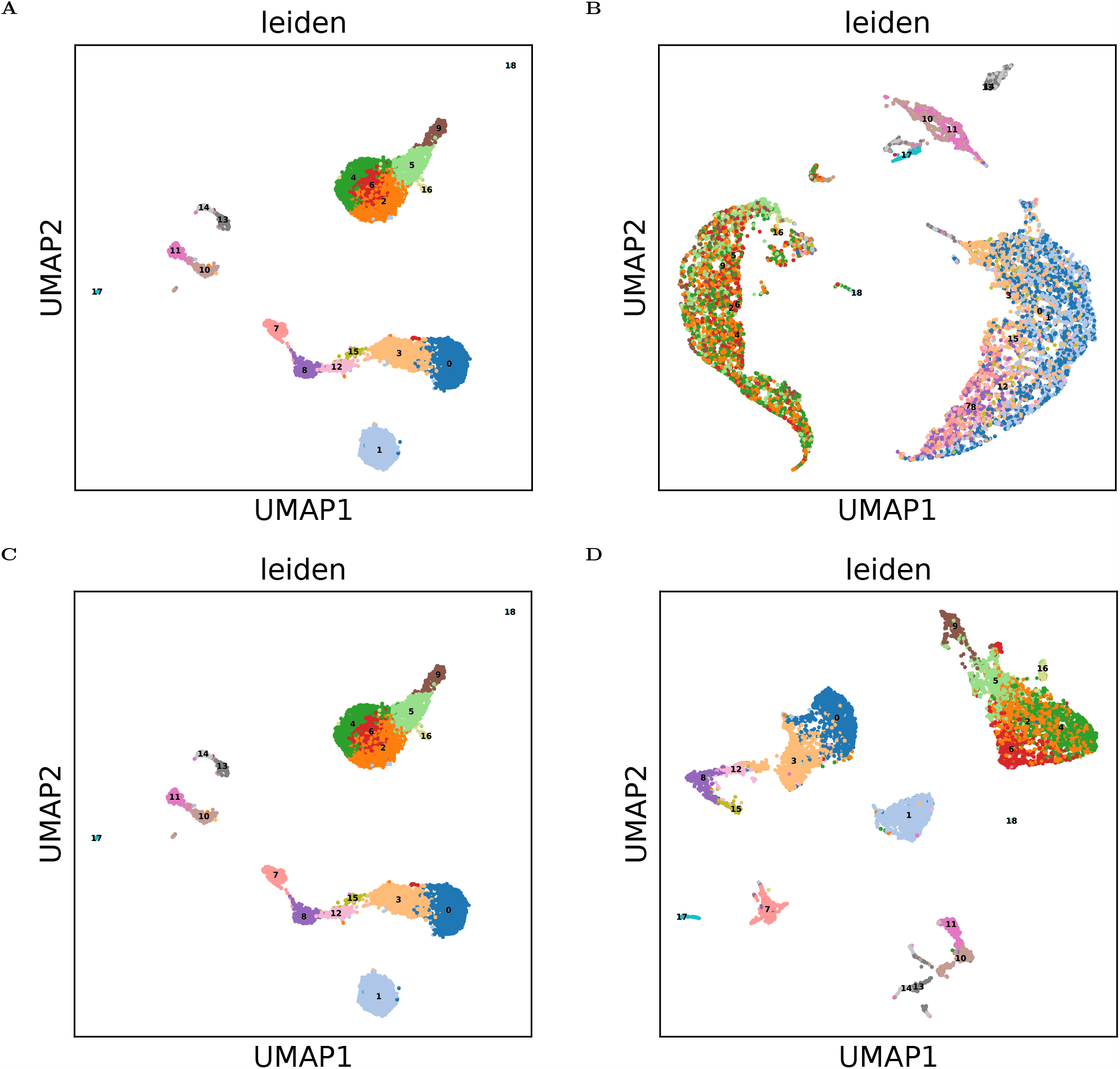
Anscombe Residuals do not require value clipping. UMAP visualization, using default MUFFIN cluster labels of the PCA representation obtained after : **A**. Anscombe residuals transform, no value clipping. **B**. Pearson Residuals transform, no value clipping. **C**. Anscombe residuals transform, clipped to +-sqrt(n) (n being the number of observations/cells, SCTransform V1 default). **D**. Pearson Residuals, clipped to +-sqrt(n). **E**. Anscombe residuals transform, clipped to +-sqrt(n/30) (SCTransform V2 default). **F**. Pearson Residuals, clipped to +-sqrt(n/30). **G**. Anscombe residuals transform, clipped to +-3. **H**. Pearson Residuals, clipped to +-3. Large clipping values can produce incoherent clusters with pearson residuals and low values can eliminate biological signal with both methods, especially in the B-Cell clusters (10,11,13,14).

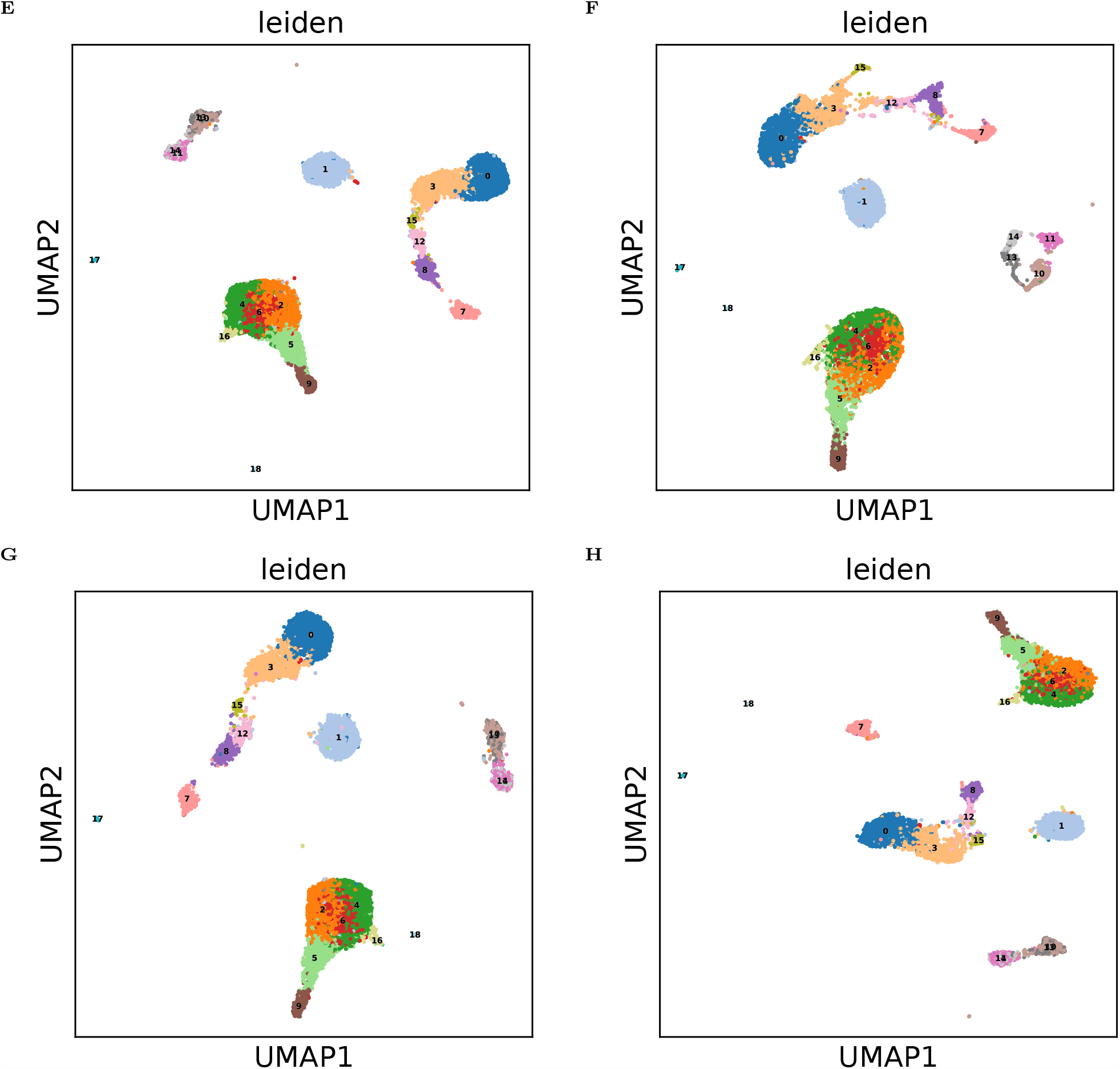

**Fig. S3.**
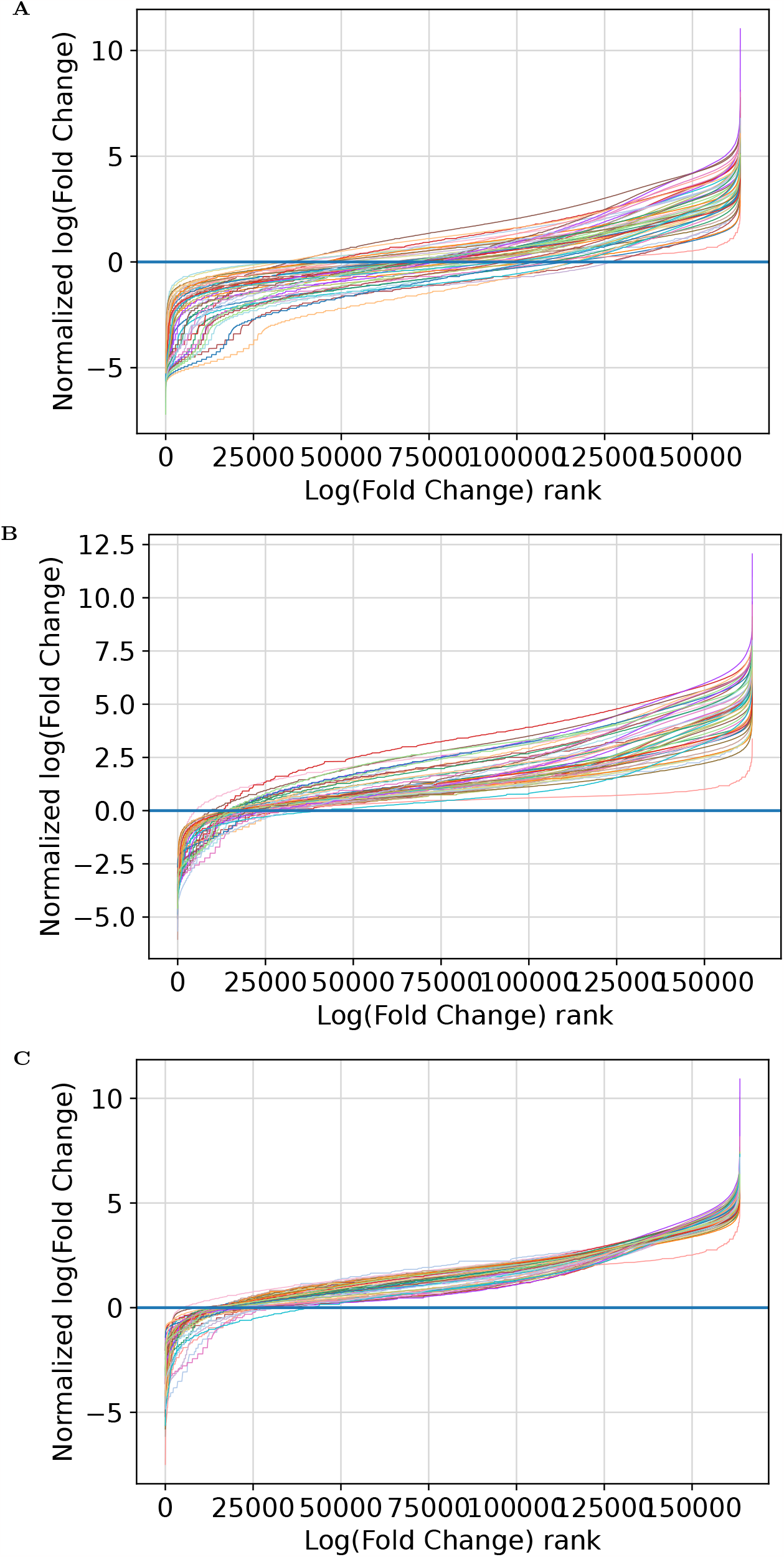
Center-and-scale normalization for input-based sequencing data. **A**. Ranked, non normalized log fold change over input curves. Each curve correspond to one experiment in the H3K4Me3 ENCODE Immune cell atlas. **B**. Ranked, centered log fold change over input curves. **C**. Ranked, centered and scaled log fold change over input curves.

**Fig. S4.**
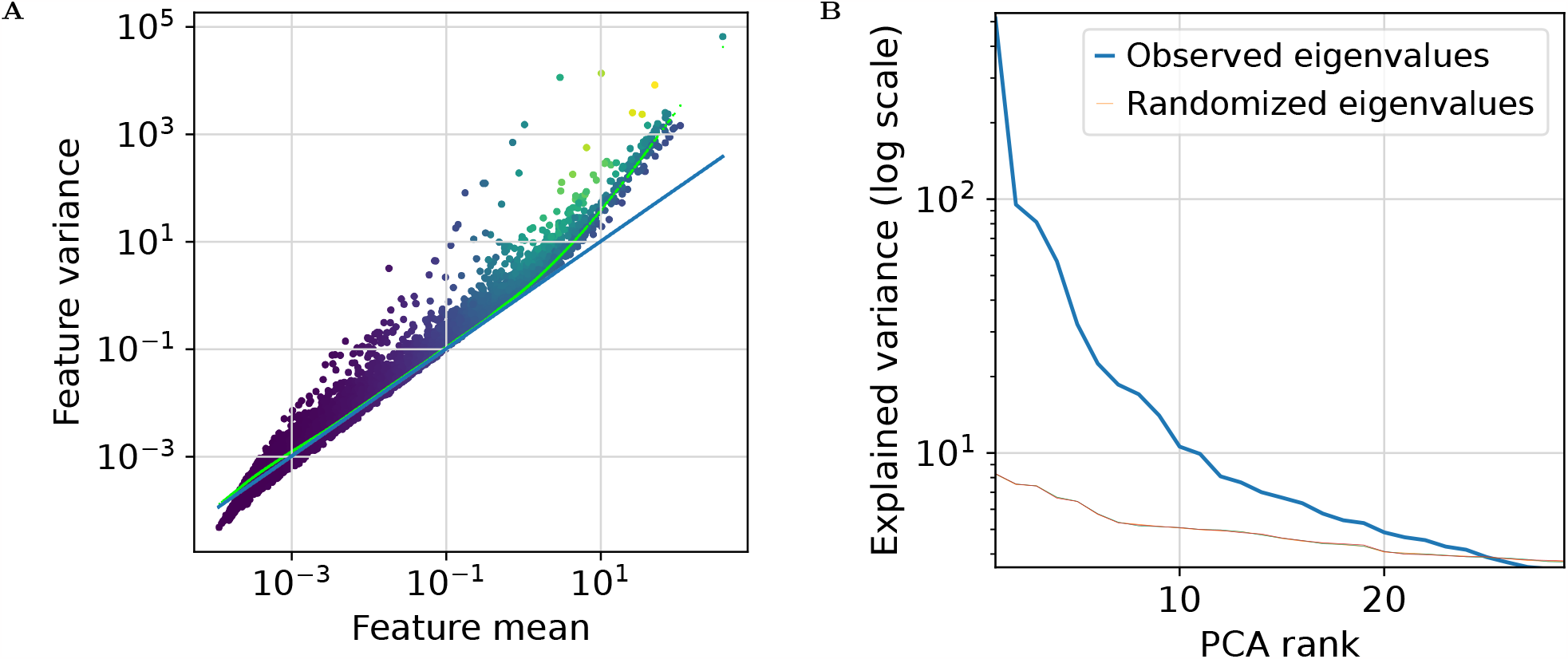
Feature weighting of Anscombe residuals, Permutation PCA. In the 10k PBMCs single-cell dataset : **A**. Mean-variance relationship of gene-expression (scran-normalized UMIs). Blue line indicates a Poisson meanvariance relationship, Green line indicates fitted mean-variance relationship. The color of each dot corresponds to its standard deviation (Yellow = larger). A lower standard deviation will result in less weight being given to the feature in downstream tasks. **B**. Observed and expected variance in each PCA component. Here 3 permutation steps have been performed (stacked Randomized eigenvalues curves).

